# Improved virus isoelectric point estimation by exclusion of known and predicted genome-binding regions

**DOI:** 10.1101/2020.07.13.201764

**Authors:** Joe Heffron, Brooke K. Mayer

**Affiliations:** Department of Civil, Construction and Environmental Engineering, Marquette University, 1637 W. Wisconsin Ave., Milwaukee, WI 53233

**Keywords:** Capsid, DNA-binding, electrostatic, modeling, point of zero charge, polynucleotide, RNA-binding

## Abstract

Accurate prediction of the isoelectric point (pI) of viruses is beneficial for modeling virus behavior in environmental transport and physical/chemical treatment applications. However, the empirically measured pIs of many viruses have thus far defied simple explanation, let alone prediction, based on the ionizable amino acid composition of the virus capsid. Here, we suggest an approach for predicting virus pI by excluding capsid regions that stabilize the virus polynucleotide via electrostatic interactions. This method was applied first to viruses with known polynucleotide-binding regions (PBRs) and/or 3D structures. Then, PBRs were predicted in a group of 32 unique viral capsid proteome sequences via conserved structures and sequence motifs. Removing predicted PBRs resulted in a significantly better fit to empirical pI values. After modification, mean differences between theoretical and empirical pI values were reduced from 2.1 ± 2.4 to 0.1 ± 1.7 pH units.

**Importance:** This model is the first to fit predicted pIs to empirical values for a diverse set of viruses. The results suggest that many previously-reported discrepancies between theoretical and empirical virus pIs can be explained by coulombic neutralization of PBRs of the inner capsid. Given the diversity of virus capsid structures, this nonarbitrary, heuristic approach to predicting virus pI offers an effective alternative to a simplistic, one-size-fits-all charge model of the virion. The accurate, structure-based prediction of PBRs of the virus capsid employed here may also be of general interest to structural virologists.

## 1. Introduction

Electrostatic interactions between virus particles and their environment are integral to virus fate and transport in physical/chemical processes and the natural environment. Virus surface charge varies between net negative and positive charges with increasing pH. A virus’ isoelectric point (pI) is defined as the pH at which the net virion charge is neutral. Knowing the pI of a virus enables prediction of virus surface charge in the environment. Predicting virus surface charge is important not only for enhancing understanding of virus deposition on surfaces such as soil particles, or virus destabilization via coagulants in water treatment, but also virion-virion aggregation (1, 2). Viruses tend to aggregate near the pI due to negated electrostatic repulsion, and aggregation can significantly impair the efficacy of disinfection processes (3, 4). In addition, knowledge of virus pI can inform virus concentration and detection in environmental samples, *e.g*., via isoelectric focusing (5, 6).

Though the available data for virus pIs are sparse and include some outliers, the general consistency of empirically determined virus pIs between researchers and spanning decades is encouraging. In their indispensable review of virus pIs, Michen and Graule (7) note that the range of reported pIs for bacteriophages MS2 and ΦX174 can be limited to 0.8 and 0.1 pH units, respectively, by limiting for strain and purity. Empirical pIs are also similar between strains of a single virus species, as shown in Figure 1. The similarity of pI between closely related viruses suggests that pI may be predictable based on conserved virion structure.

**Figure 1:**
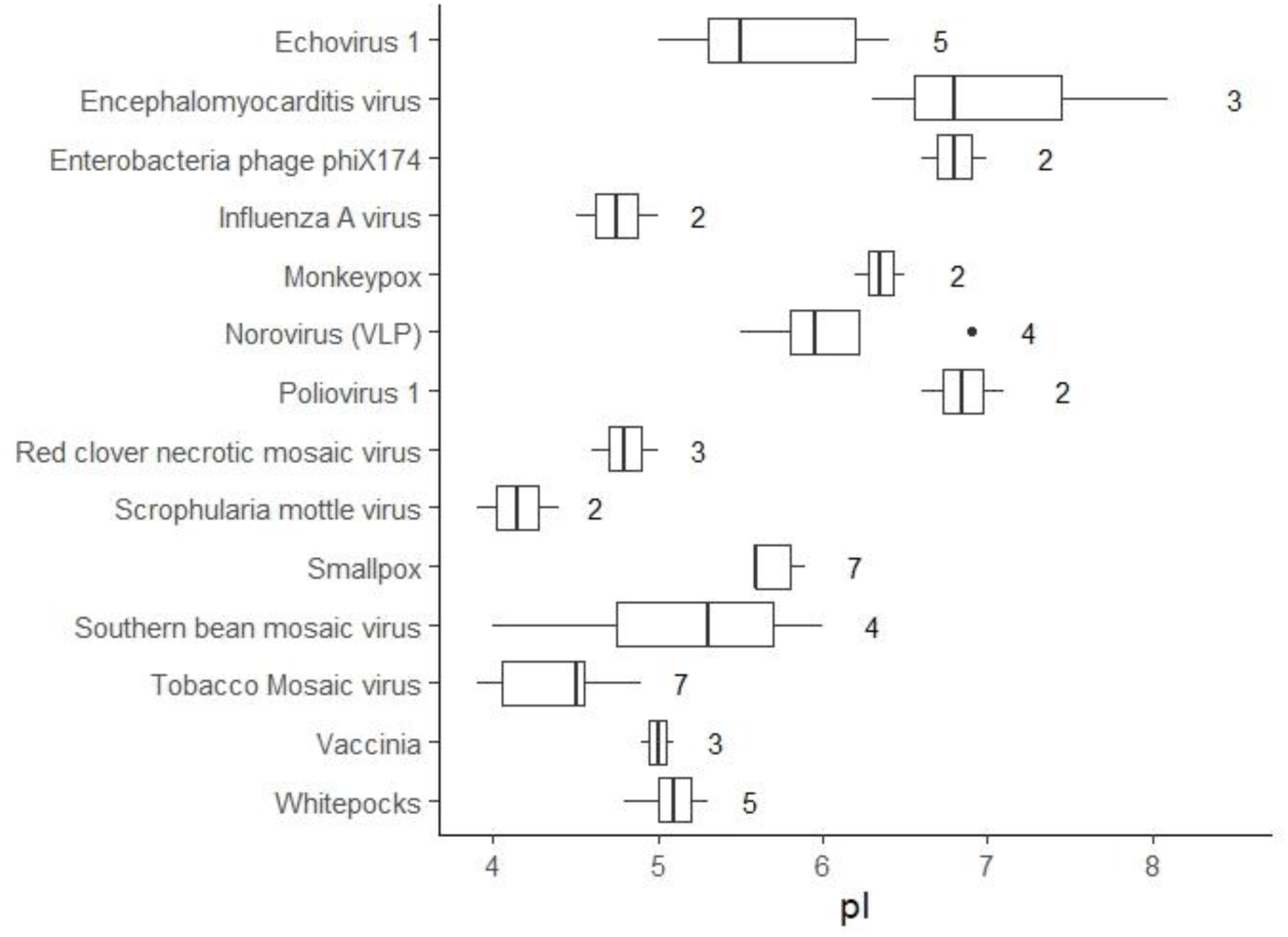
Relative conservation of isoelectric point (pI) in closely related virus strains. The plot shows empirical pI values for different strains of the same species, with the number of strains shown to the right of each box (data summarized from Michen and Graule (7)). To minimize differences due to experimental conditions, pI values for each virus along the y-axis were obtained from single studies comparing different strains.

Attempts to model virion pI generally involve quantifying and modifying the charges of ionizable amino acids within capsid proteins (8–12). Altering the composition of ionizable amino acids within capsid proteins can have a predictable and measurable effect on virion pI (13). Modern recombinant techniques allow some degree of “charge tuning” of viral particles by adding or replacing ionizable amino acids within capsid proteins (14).

However, the sequence of ionizable amino acids alone appears insufficient to accurately predict pI. Based on analyses of hundreds of virus proteomes, theoretical virus pIs calculated from ionizable amino acid residues are tightly clustered near neutral pH, with an overall range between approximately pH 5.5 and 8 (15, 16). However, empirical virus pIs are frequently reported below pH 5, and have been measured as low as pH 2 (7). Given that the distribution of theoretical virus pIs (pH 5.5 – 8) is on the same order as the variation in empirical pIs between strains of a single virus species (< 2 pH units, see Figure 1), refined prediction of virus pI based on ionizable amino acids may not be possible at the species level. Rather, the primary goal for a model of virion pI should be to reliably predict which viruses will have pIs outside the expected circumneutral range.

Previous researchers have explained the differences between theoretical and empirical pIs by either supposing a strong, negative influence from the viral polynucleotide (genome) at the virion core (12, 17–21), or by supposing that only the exterior capsid surface contributes to virion charge (9, 11). Given the low pKa (∼1) of the polynucleotide phosphodiester group (Table 1) and the porous nature of virus capsids, the assumption that the virion core influences overall charge is credible. However, DNA- and RNA-folding and compaction during encapsidation requires a cloud of counterions to overcome electrostatic self-repulsion (22–26), at least some of which are likely retained in the assembled virion core (27). In addition, experiments comparing the charge of whole virions to empty capsids lacking a genome (virus like particles) have failed to account for major discrepancies in theoretical and empirical pIs (9, 19, 28, 29). The second proposed model, in which only exterior capsid residues contribute to virion charge, has been used to calculate predicted pIs, but only for structurally similar bacteriophages in the *Leviviridae* family (9, 11). Neither of these previous methods (presuming a negative charge from the virion core or selecting only exterior residues) has yet been demonstrated on a large, diverse set of viruses.

**Table 1:** Acid dissociation constants (pKas) for protein and nucleic acid constituents.

|  | Residue | pKa | Charge |
| --- | --- | --- | --- |
| **Protein\*** |  |  |  |
|  | Terminal carboxyl | 2.87 | Negative |
|  | Aspartic acid | 3.87 | Negative |
|  | Glutamic acid | 4.41 | Negative |
|  | Cysteine | 7.56 | Negative |
|  | Tyrosine | 10.85 | Negative |
|  | Histidine | 5.64 | Positive |
|  | Lysine | 9.05 | Positive |
|  | Terminal amine | 9.09 | Positive |
|  | Arginine | 11.84 | Positive |
| **Nucleic Acid\*\*** |  |  |  |
|  | Primary phosphoryl | 1 | Negative |
|  | Deprotonated base (G, T, U) | 9.4 - 10 | Negative |
|  | Amino group (A, C, G) | 2.3 - 4.6 | Positive |

\* values from Isoelectric Point Calculator (isoelectric.org) (71)
\*\* values from Blackburn, 2006 (72)

Furthermore, using either approach to develop a predictive model of pI would require fitting one or more variables to the empirical pI data, since there is no other empirical source of data to describe the influence of core charge or decision criterion for what constitutes an “exterior” residue in large and convoluted capsids. Given the limited empirical pI data and bias toward viruses commonly used in research (*e.g*., *Leviviridae* and plant viruses), these approaches are likely to overfit a prediction to the available data. Preferably, a model for pI prediction would rely on a separate, independently verifiable criterion for what elements of virion structure contribute to the overall charge.

The goal of this study was to propose a simple model to improve virus pI prediction. Our hypothesis was that positive charges from basic protein residues in polynucleotide-binding regions (PBRs) of the capsid interior are neutralized by noncovalent bonding with the viral polynucleotide (genome), and therefore should not be considered in the capsid pI calculation. We approached this challenge by first modifying capsid protein sequences based on regions known or suspected to stabilize the viral genome, and then calculated the predicted pI using a simple sum of charges method. This heuristic approach applies a rule for including and excluding amino acids from the pI calculation, rather than imposing a simplified physical model on the virion structure. In addition, the heuristic is nonarbitrary, in that amino acids are excluded based on function, rather than an attempt to fit the predicted pIs to empirical values. For this study, both 3D structural models of virus capsids and capsid proteome sequences were evaluated with and without modifications. The implication of this approach is that while one simple structural model may not be applicable to all viruses, a descriptive model of virion pI can arise from a simple, nonarbitrary heuristic.

## 2. Results

The evidence in support of PBR exclusion as a means of pI prediction comes from both known capsid structures and predicted PBRs. “Known structures” include both 3D capsid models and experimentally-determined PBRs. Given the relatively few viruses for which both known structures and empirical pI values were available, PBRs were predicted for a larger set of viruses based on conserved structures apparent in the experimentally-determined PBRs. The most accurate PBR predictions were then used to predict pIs for the extended virus set.

### 2.1. Known capsid structures

Some researchers have suggested that only residues on the exterior capsid surfaces contribute to overall capsid charge (9, 11). However, for the set of viruses with available 3D structures and empirical pIs, residues on the exterior surface were a poor predictor of overall virion charge. As shown in Figure 2, exterior surface residues identified via CapsidMaps (30) tended to be composed of only acidic or basic residues, with no correlation to empirical pI. Interestingly, removing only the residues on the interior capsid surface did result in a better fit between theoretical and empirical pIs compared to unmodified capsids (Figure 2). Many virus capsids feature a concentration of basic residues toward the interior – notably the ssRNA phages of *Leviviridae* commonly used in research. Thus, the exterior residues of these viruses have a lower pI than the capsid as a whole. However, whether a given virus has this unbalanced distribution of ionizable amino acids has defied any straightforward prediction (10). Given the diverse virus sizes and morphologies represented in Figure 2, a single charge distribution model is unlikely to explain why only the innermost surface-exposed residues do not contribute to overall virion charge.

**Figure 2:**
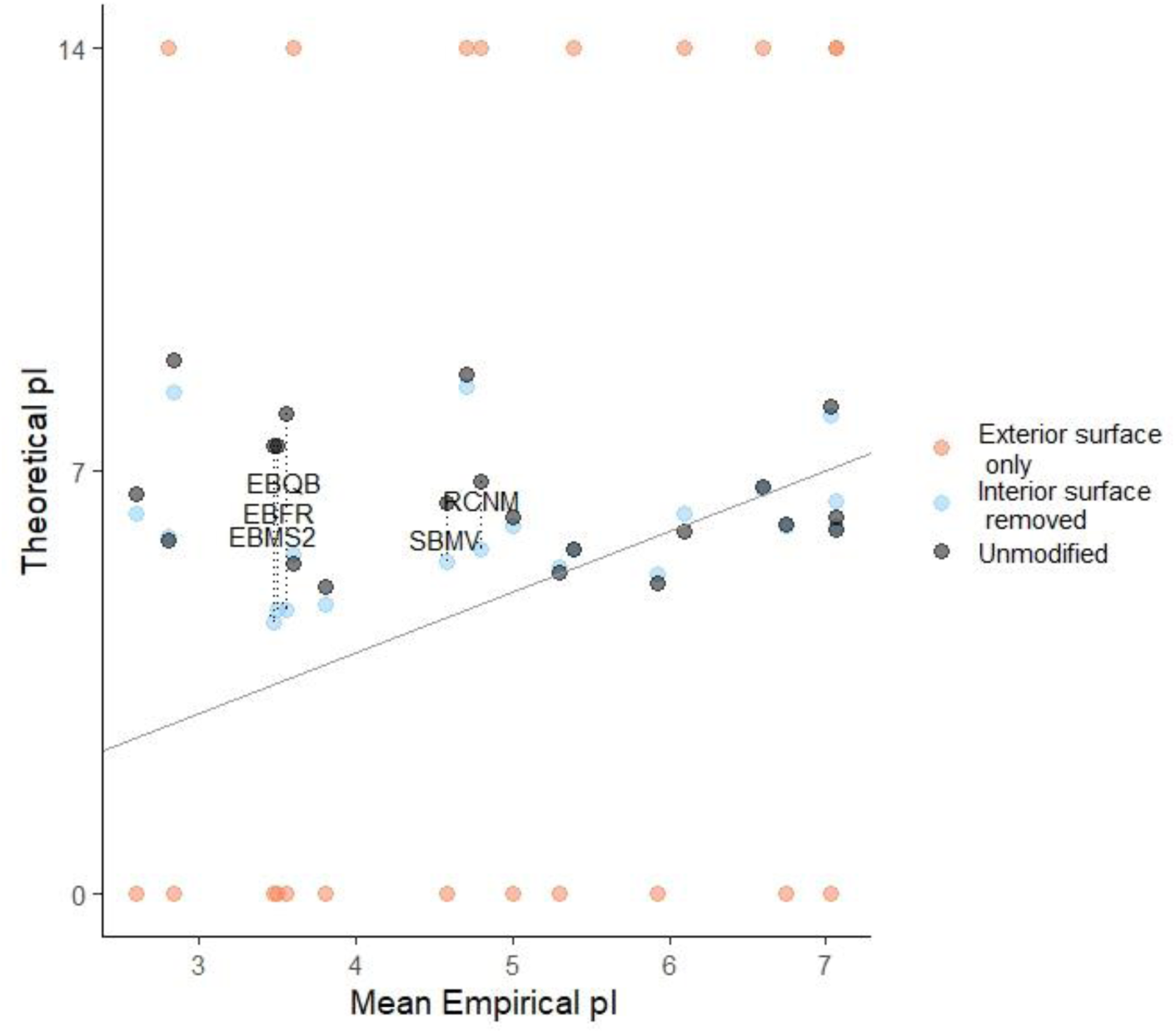
Impact of including only exterior residues in predicted pI calculation using 3D capsid structures. Exterior residues are defined by inclusion of only the exterior capsid surface or exclusion of the interior capsid surface, as identified via CapsidMaps (30). Exterior capsid surfaces were composed of entirely acidic or basic residues; these pIs are shown as approaching neutral charge at pH 0 or 14, respectively. The mean empirical pI value for each unique virus is shown in comparison to the virus’ predicted pI calculated from exterior capsid residues, as defined in the legends. The diagonal line represents equivalent theoretical and empirical pIs. Viruses showing the greatest improvement in predicted pI after removing interior surfaces are labeled; a guide to virus abbreviations is provided in Table 2.

A review of the viruses most positively impacted by removal of interior residues provides some insight. The five viruses with the greatest improvement in pI estimation are labeled in Figure 2: *Leviviridae* bacteriophages fr (EBFR), MS2 (EBMS2), and Qβ (EBQB); as well as ssRNA plant viruses red clover necrotic mosaic virus (RCNM), and southern bean mosaic virus (SBMV). *Leviviridae* feature a highly-conserved assembly mechanism by which RNA binds to planar beta sheets on the capsid protein interior, forming subunits from which the capsid self-assembles (31– 34). Thus, these phages have large, basic surfaces on the capsid interior devoted to RNA binding. The other two viruses, SBMV and RCNM, are ssRNA plant virus with highly basic, disordered N-termini. These disordered regions also function to stabilize the viral polynucleotide (35–37). Therefore, the viruses showing the greatest impact from interior residue exclusion all feature highly basic interior residues that stabilize the viral genome. Both of these basic, interior capsid features are noncovalently bound to the viral RNA (31, 33, 35, 36), and therefore their positive contribution to virion charge is likely negated by the negatively charged polynucleotide.

**Table 2:** Classification and abbreviations for viruses used in this study.

| Abbrev. | Species (Strain) | NCBI<br>Taxon* | PDBID<sup>†</sup> | Genus | Family | Nucleic<br>Acid | Host<br>Kingdom |
| --- | --- | --- | --- | --- | --- | --- | --- |
| AAV4 | Adeno-associated virus 4 | 57579 | 2g8g (73) | Dependo-<br>parvovirus | Parvoviridae | ssDNA | Animalia |
| BDMV | Belladonna mottle virus | 12149 | -- | Tymovirus | Tymoviridae | ssRNA | Plantae |
| BP29 | *Bacillus* phage Φ29 | 10756 | -- | Φ29virus | Podoviridae | dsDNA | Bacteria |
| CCMV | Cowpea chlorotic mottle<br>virus | 12303 | 1cwp (74) | Bromovirus | Bromoviridae | ssRNA | Plantae |
| CMV | Cucumber mosaic virus<br>(FNY) | 12307 | 1f15 (75) | Cucumovirus | Bromoviridae | ssRNA | Plantae |
| CPaV2 | Canine parvovirus 2 | 10790 | -- | Proto-<br>parvovirus | Parvoviridae | ssDNA | Animalia |
| CPaV2** | Feline panleukopenia<br>virus | 10787 | 1c8g (76) | Proto-<br>parvovirus | Parvoviridae | ssDNA | Animalia |
| CRPV | Cottontail rabbit<br>papillomavirus (Kansas) | 31553 | -- | Kappa-<br>papillomavirus | Papilloma-<br>viridae | dsDNA | Animalia |
| CRPV** | Human papillomavirus 16 | 333760 | 5keq (77) | Alpha-<br>papillomavirus | Papilloma-<br>viridae | dsDNA | Animalia |
| CXA21 | Coxsackievirus A21 | 12070 | 1z7s (78) | Enterovirus | Picornaviridae | ssRNA | Animalia |
| CXB5 | Human coxsackievirus B5<br>(Peterborough) | 103907 | -- | Enterovirus | Picornaviridae | ssRNA | Animalia |
| CXB5** | Human coxsackievirus B3 | 103904 | 1cov (79) | Enterovirus | Picornaviridae | ssRNA | Animalia |
| EBFR | Enterobacteria phage fr | 12017 | 1frs (80) | Levivirus | Leviviridae | ssRNA | Bacteria |
| EBGA | Enterobacteria phage GA | 12018 | 1gav (34) | Levivirus | Leviviridae | ssRNA | Bacteria |
| EBMS2 | Enterobacteria phage MS2 | 329852 | 2ms2 (81) | Levivirus | Leviviridae | ssRNA | Bacteria |
| EBQB | Enterobacteria phage Qβ | 39803 | 5vly (82) | Allolevivirus | Leviviridae | ssRNA | Bacteria |
| EBSP | Enterobacteria phage SP | 12027 | -- | Allolevivirus | Leviviridae | ssRNA | Bacteria |
| EBT4 | Enterobacteria phage T4 | 10665 | 5vf3 (83) | T4likevirus | Myoviridae | dsDNA | Bacteria |
| ECV1 | Echovirus 1 (Farouk) | 103908 | 1ev1 (84) | Enterovirus | Picornaviridae | ssRNA | Animalia |
| ELV | Erysimum latent virus | 12152 | -- | Tymovirus | Tymoviridae | ssRNA | Plantae |
| HAdV5 | Human adenovirus 5 | 28285 | 4v4u (85) | Mast-adenovirus | Adenoviridae | dsDNA | Animalia |
| HHAV | Hepatitis A virus (HM175) | 12098 | 4qpi (86) | Hepatovirus | Picornaviridae | ssRNA | Animalia |
| HRV2 | Human rhinovirus 2 | 12130 | 1fpn (87) | Enterovirus | Picornaviridae | ssRNA | Animalia |
| MEV | Mengo encephalomyocarditis virus | 12107 | 2mev (88) | Cardiovirus | Picornaviridae | ssRNA | Animalia |
| NOR1 | Norwalk Virus (Funabashi) | 524364 | 1ihm (89) | Norovirus | Caliciviridae | ssRNA | Animalia |
| PHIX | Enterobacteria phage ΦX174 (Sanger) | 10847 | 2bpa (90) | ΦX174-microvirus | Microviridae | ssDNA | Bacteria |
| PM2 | Pseudoalteromonas phage PM2 | 10661 | 2w0c (91) | Corticovirus | Corticoviridae | dsDNA | Bacteria |
| POL1 | Poliovirus (Mahoney) | 12081 | 1hxs (92) | Enterovirus | Picornaviridae | ssRNA | Animalia |
| PRD1 | Enterobacteria phage PRD1 | 10658 | 1w8x (93) | Alphatectivirus | Tectiviridae | dsDNA | Bacteria |
| RCNM | Red clover necrotic mosaic virus | 12267 | 6mrm (94) | Dianthovirus | Tombusviridae | ssRNA | Plantae |
| REO3 | Reovirus 3 (Dearing) | 10886 | 2cse (95) | Orthoreovirus | Reoviridae | dsRNA | Animalia |
| SBMV | Southern bean mosaic virus | 652938 | 4sbv (96) | Sobemovirus | Solemoviridae | ssRNA | Plantae |
| ScrMV | Scrophularia mottle virus | 312273 | -- | Tymovirus | Tymoviridae | ssRNA | Plantae |
| SRVA | Simian rotavirus A (SA11) | 450149 | 4v7q (97) | Rotavirus | Reoviridae | dsRNA | Animalia |
| TBMV | Tobacco mosaic virus (Vulgare) | 12243 | -- | Tobamovirus | Virgaviridae | ssRNA | Plantae |
| TYMV | Turnip yellow mosaic virus | 12154 | 1auy (98) | Tymovirus | Tymoviridae | ssRNA | Plantae |

\* NCBI Taxon: National Center for Biotechnology Information (www.ncbi.nlm.nih.gov) taxonomical ID (99)
† PDBID: Protein Data Bank (rcsb.org) ID used for 3D structural comparisons (100)
\*\* Alternate species/strain used for 3D structure only

### 2.2. Known polynucleotide-binding regions

To determine the impact of PBRs on virus pI, known PBRs were identified for 15 viruses from annotations in the UniProt database (38) and the literature (see Table S3). Theoretical virus pIs were compared before and after modification by excluding PBRs from the charge calculation. As shown in Figure 3, excluding PBRs improved pI estimation for the majority of viruses. The predicted pIs before modification deviated from empirical values by 2.2 ± 2.4 pH units; after modification, predicted pIs deviated from empirical pIs by 1.4 ± 1.5. This reduction in mean deviation after PBR exclusion was significant to a high degree of confidence (paired t = 5.81 (14 df), p = 5×10^−5^). By comparison, excluding interior surface residues also decreased the deviation between theoretical and empirical pIs to 1.0 ± 1.7 pH units, though the improvement was slightly less significant (paired t = 2.82 (20 df), p = 0.01). However, the identification of capsid surfaces may not be practical for larger, layered capsids due to complex structure, computational burden or lack of entire 3D structures.

**Figure 3:**
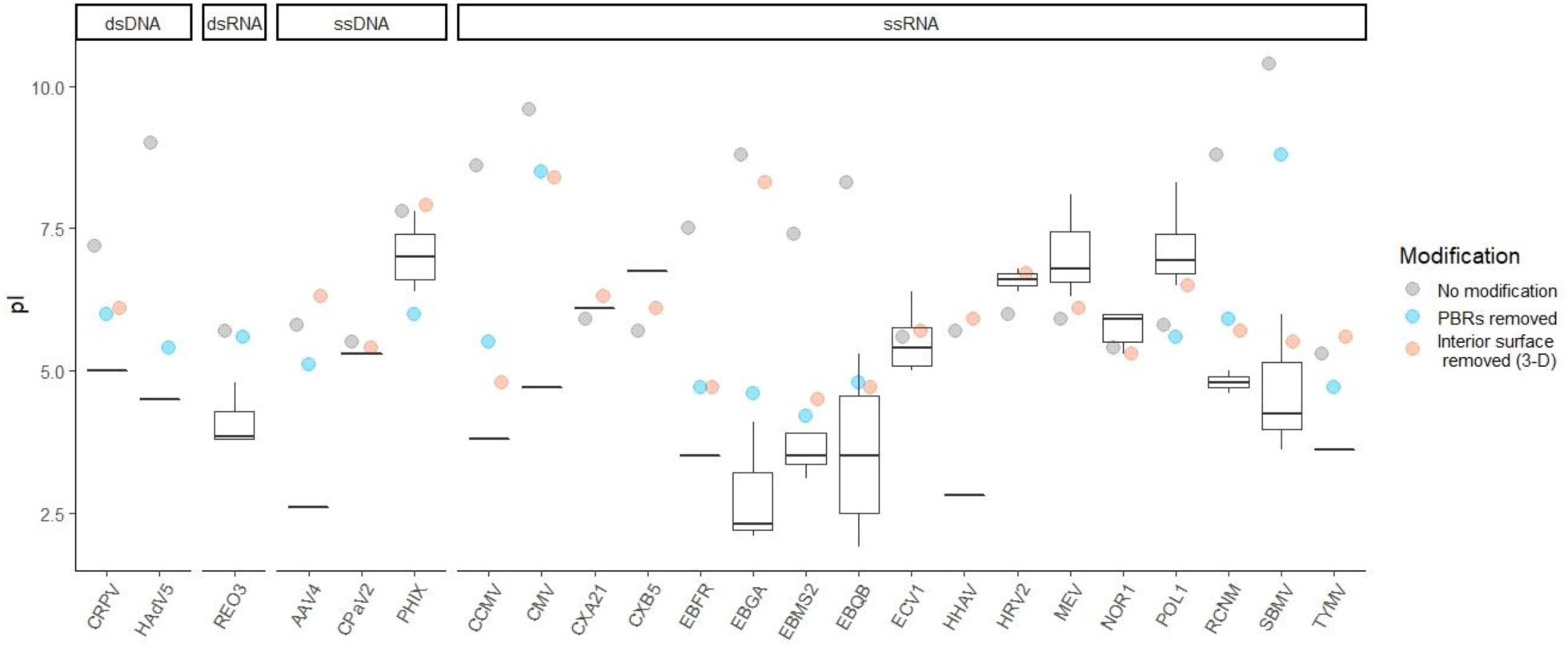
Effect of modifications on theoretical capsid pI. Box and whisker plots represent the range of empirical pIs found in the literature, while circles represent predicted pIs. The predicted pIs reflect the theoretical capsid charges: without modification, after excluding known viral polynucleotide-binding regions (“PBRs removed”) and after removing capsid interior surfaces using 3D structures (“Interior surface removed (3D)”). Both modifications were not possible for some viruses due to unknown PBR locations, unavailable 3D structures, and/or large size. A key to the 23 virus abbreviations (x axis) is provided in Table 2.

Modification via PBR exclusion and interior surface exclusion appeared to be complementary. Many viruses in Figure 3 lacked either known PBRs or capsid structures, so only one method was possible with the available data. In only a few cases, either PBR or interior surface exclusion produced a far better prediction than the other method. For example, pI estimation for bacteriophage GA (EBGA) was far closer to empirical values via PBR exclusion. The interior of the bacteriophage GA capsid protein contains both acidic and basic residues (34), so non-specific removal of the entire interior surface has little net effect on capsid pI. On the other hand, southern bean mosaic virus (SBMV) was more accufrately predicted after interior surface exclusion, possibly indicating that the full extent of the PBR was not known for this virus.

In this study, PBRs with the greatest impact on predicted pI could be divided into three primary categories: predominately basic beta sheets and associated turns, disordered polypeptide termini (primarily N-termini), and histone-like proteins. Basic beta sheets were typical PBRs for the *Leviviridae* family of ssRNA bacteriophages routinely used as model viruses in research. Some *Leviviridae* phages (fr, MS2 and SP) also had basic alpha helices serving as PBRs in their maturation and minor capsid proteins. In the two representatives of *Allolevivirus* (bacteriophages Qβ and SP), basic residues occurred primarily on the turns between beta sheets on the major capsid protein, rather than on the beta sheets themselves. Four other ssRNA viruses in this study – cowpea chlorotic mosaic virus (CCMV), cucumber mosaic virus (CMV), red clover necrotic mosaic virus (RCNM), and southern bean mosaic virus (SBMV) – featured PBRs within disordered, highly basic N-termini (35, 36). Some of these N-termini also feature alpha helices that bind to RNA stem-loops (39). Human adenovirus C5 (HAdV5) was the only virus in this study that featured histone-like proteins, and the removal of these proteins improved the accuracy of the pI prediction by 3.6 pH units, as shown in Figure 3. Reovirus type 3 (REO3) also has a protein thought to act as a spool for RNA within the capsid (40); however, inclusion or exclusion of this protein did not impact the predicted pI.

In addition to common PBR structures, PBR sequences in this study had high arginine and lysine fractions (0.42 ± 0.29). These arginine- and lysine-rich regions are often indicative of RNA- and DNA-binding (41–44). The beta sheets of *Leviviridae* capsid proteins had lower arginine/lysine fractions (0.11 ± 0.01) than other PBRs, as basic amino acids tend to be distributed over a non-contiguous surface rather than within one short sequence. A full list of viral proteins and their PBRs is provided in the Supporting Information (Table S3).The set of viruses evaluated in Figure 3 was limited by the availability of empirical pIs, as well as curated proteome sequences and 3D structures. Here, as in other virus pI research, *Leviviridae* are in particular over-represented, preempting the conclusion that the PBR exclusion approach is universally applicable. Nonetheless, excluding PBRs explained multiple discrepancies in predicted pIs with a single, nonarbitrary heuristic, and was thus a promising direction for a predictive model of virus pI. However, PBRs would have to be predicted within virus proteome sequences in order to apply this method to a larger set of viruses.

### 2.3. Predicted polynucleotide-binding regions

Unfortunately, excluding PBRs by the above methods required either a high-resolution 3D capsid model or the full extent of the capsid PBR(s). A method of predicting PBRs based on capsid protein sequences would be far preferable, as capsid proteomes are available for a wide range of viruses. Furthermore, PBRs are often discovered by point mutations/deletions of select amino acids, so the full extent of the PBR may not be known. As a first attempt at a predictive model of virion pI based on PBR exclusion, PBRs were predicted in a diverse group of 32 viruses based on the conserved PBR features discussed in Section 2.2. Specifically, PBR predictions attempted to capture basic beta sheets and associated beta turns, disordered polypeptide termini, and arginine-rich regions. (Though both arginine and lysine contributed to the predicted PBRs, the common term “arginine-rich” is used here to describe these basic regions, since arginine is the dominant residue.) In addition, two web-based tools for detecting XNA-binding residues, Pprint (45, 46) and DRNApred (47, 48), were evaluated for virus PBR prediction. Both tools modeled the likelihood of amino acids binding to RNA or DNA based on position within the primary sequence. All PBR predictions were evaluated against the known PBRs discussed in Section 2.2, as well as a validation set of 40 other capsid proteins. Predicted pIs were then calculated by excluding the predicted PBRs; predicted pIs were compared to empirical values.

#### 2.3.1. Prediction of polynucleotide-binding regions

Overall, searching for conserved structures offered the most reliable PBR prediction. As a single predictor, arginine-rich regions had greatest predictive power (MCC = 0.32 ± 0.29). Prediction of PBRs via beta structures (sheets: MCC = 0.05 ± 0.23, turns: MCC = 0.07 ± 0.20) and disordered termini (MCC = 0.16 ± 0.31) had lower MCCs than the arginine-rich region prediction. However, these low metrics may reflect the relatively low prevalence of beta sheets and disordered termini within the training and validation sets, rather than a poor match to experimental data. These structures were intended to complement one another, rather than predict all regions with a single structure. When evaluated only on proteins containing beta sheet PBRs, the beta sheet predictor performed far better (MCC = 0.45 ± 0.16). The disordered termini also performed better against a set of only proteins with PBRs located on disordered termini (MCC = 0.35 ± 0.32), though the standard deviation indicates that the fit was not universal or specific. A summary of MCCs for all predictions is provided in the Supporting Information (Table S1).

Most combinations of structures failed to improve on the prediction via arginine-rich regions alone. PBR prediction based on arginine-rich regions alone covered 18% of residues predicted based on beta sheets, 39% of beta turns, and 26% disordered termini. Since only the combination of arginine-rich regions and beta turns provided a better fit to training and validation data, beta sheet and disordered termini predictions likely contributed more false positives than arginine-rich regions alone. Selection of arginine-rich regions also successfully identified known alpha helix PBRs in maturation proteins of *Leviviridae* phages fr (EBFR) and MS2 (EBMS2), as well as 61% of residues in the histone-like protein of human adenovirus 5 (HAdV5). All PBR predictions based on beta sheets, beta turns, disordered termini and arginine-rich regions are provided in the Supporting Information (“PBRpredictions.pdf”).

However, beta turns in particular were complementary to the arginine-rich search, likely because bases in these regions may be adjacent in the tertiary structure but distant in the primary sequence. Fittingly, a combination of arginine-rich regions and beta turns was optimal for predicting the known PBRs in this study (MCC = 0.34 ± 0.28). Arginine-rich regions alone had a slightly higher mean MCC (0.27 ± 0.32) for the validation set than the combination of arginine-rich and beta turn regions (MCC = 0.26 ± 0.29). However, the lower variance of the latter indicated fewer poor predictions. Therefore, the combination of arginine-rich and beta turn regions was used as the preferred PBR prediction method for this study.

The naïve, structure-based PBR prediction method used here performed well compared to other predictors of XNA-binding residues. For reference, MCCs for other available XNA-binding prediction tools range from approximately 0.14 to 0.23 for RNA-binding and 0.14 to 0.35 for DNA-binding (49). The two sequence-specific predictors compared here, Pprint (MCC = 0.22 ± 0.27) and DRNApred (MCC = 0.17 ± 0.23), performed within this range for virus PBRs as well. Pprint performed better than DRNApred in this study, possibly because Pprint was optimized to the PBR training set, while DRNApred used a default decision criterion. However, both tools were below the median MCC of the structure-based predictions considered here. Neither tool was designed with virus polynucleotides or capsid proteins in mind; thus, poor performance is more a reflection of the application than the tools themselves. However, the structure-based PBR prediction used here (MCC = 0.34 ± 0.28) also performed comparably to maximum reported performance of Pprint (MCC = 0.32) (50) and DRNApred (MCC = 0.31 [DNA] and 0.36 [RNA]) (48) on their intended data sets (as determined by the respective authors).

#### 2.3.2. Impact of predicted polynucleotide-binding regions on predicted isoelectric point

The impact of excluding predicted PBRs in pI calculations can be seen for individual viruses in Figure 4. Since the combination of arginine-rich regions and beta turns provided the best prediction of PBRs, the PBRs predicted under these parameters were excluded for the “structure-based prediction” of pI. The overall improvement in accuracy of the modified predictions compared to the unmodified predictions is shown in the histogram in Figure 5. The differences between theoretical and empirical pI values decreased in both magnitude and variance after modification, from 2.1 ± 2.4 to 0.1 ± 1.7 pH units. This difference was significant to a high degree of confidence (*paired t =* 7.24 (31 df), *p =* 4×10^−8^). In addition, the histogram shifted from a bimodal to a normal distribution, indicating again that the PBR exclusion method accounts for deviations in one group (viruses with PBRs) without dramatically shifting predictions for the other (viruses with covalently-bound or free polynucleotides).

**Figure 4.**
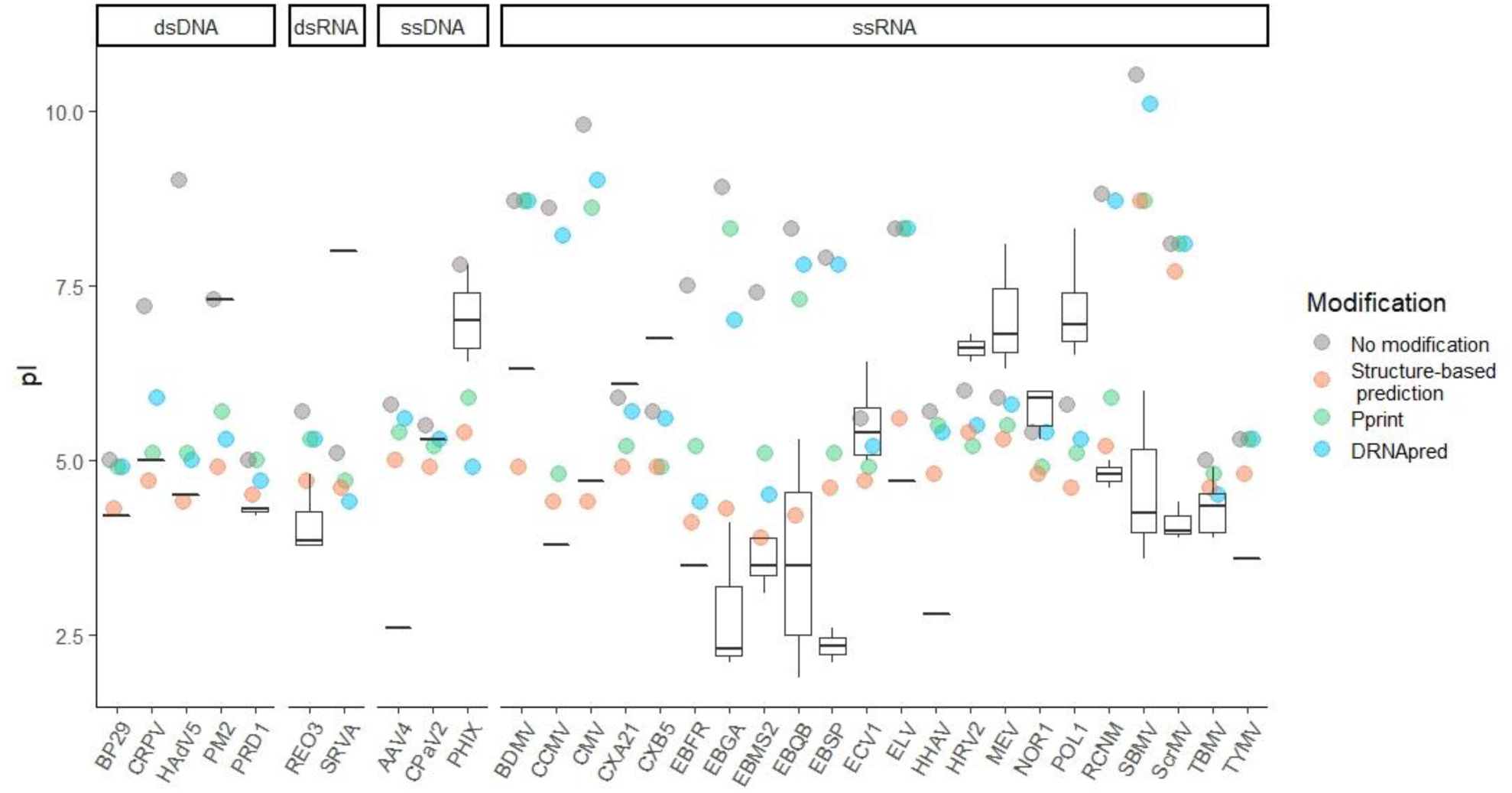
Effect of excluding predicted polynucleotide-binding regions (PBRs) on theoretical capsid pI (colored circles) for 32 viruses. Box and whisker plots represent the range of empirical pIs found in the literature, while circles represent predicted pIs calculated without modification, as well as after excluding predicted PBRs. PBRs were predicted either via the structure-based prediction developed in this study identifying arginine-rich regions and beta turns, or one of two available XNA-binding prediction tools, Pprint (45, 46) and DRNApred. A key to the virus abbreviations (x axis) is provided in Table 2.

**Figure 5.**
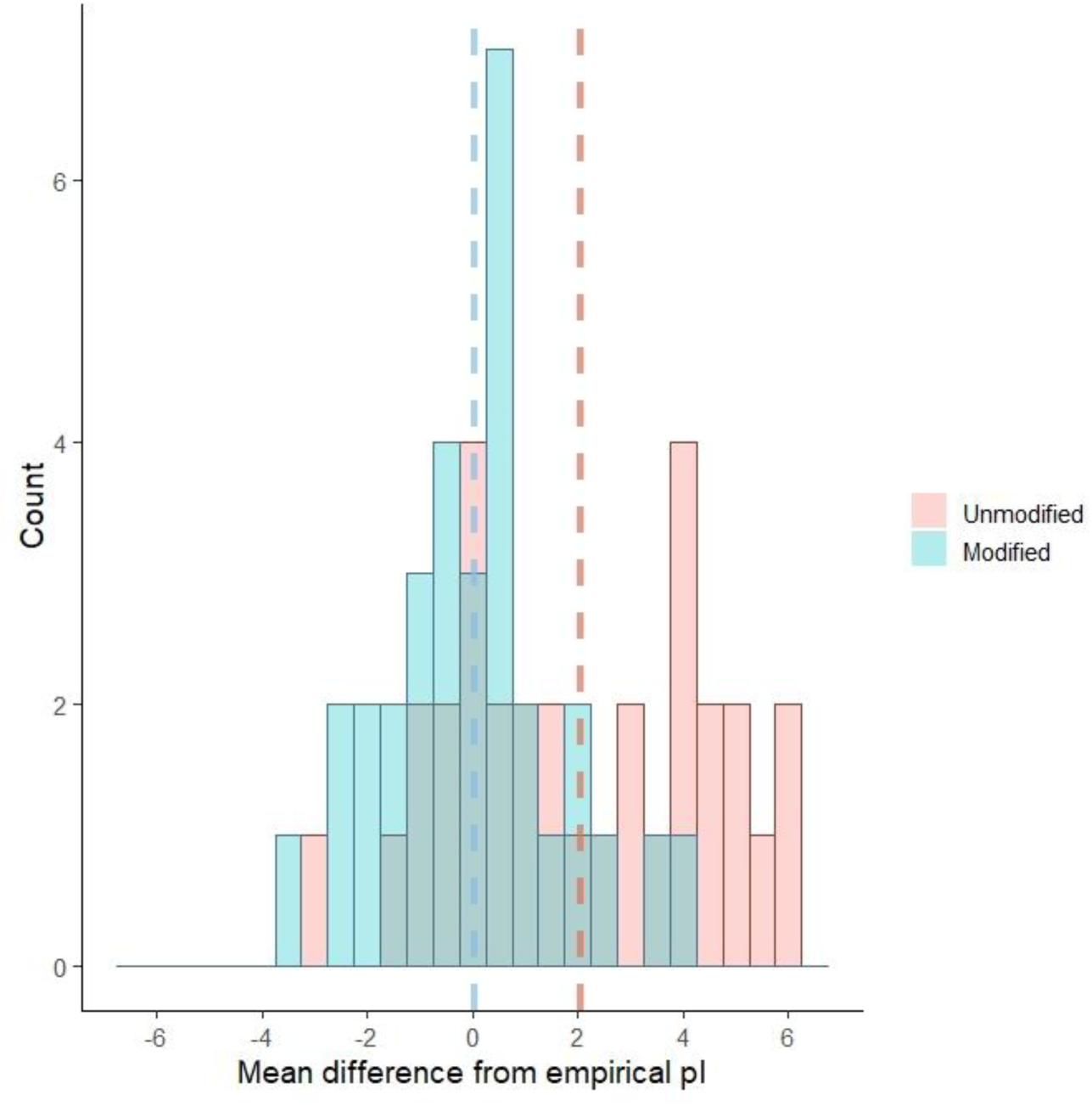
Histogram showing the shift in difference between predicted and mean empirical pI with and without modification by removal of predicted polynucleotide-binding regions (PBRs, including both arginine-rich regions and beta turns). Dashed lines represent the mean of means for each category (Modified or Unmodified), by color. The differences between theoretical and empirical pI values decreased significantly after modification, from 2.1 ± 2.4 to 0.1 ± 1.7 pH units.

Perhaps more impressively, accurate pI estimation was strongly correlated with the mean MCC of PBR prediction, as demonstrated in Figure 6. MCCs for all PBR predictions used in this study negatively correlated to the mean absolute difference between theoretical and empirical pI (Spearman’s correlation ρ = -0.84, p < 2 × 10^−16^). Though training of the PBR prediction tools occurred independently of any impact on virus pI, better PBR prediction resulted in better pI prediction.

**Figure 6.**
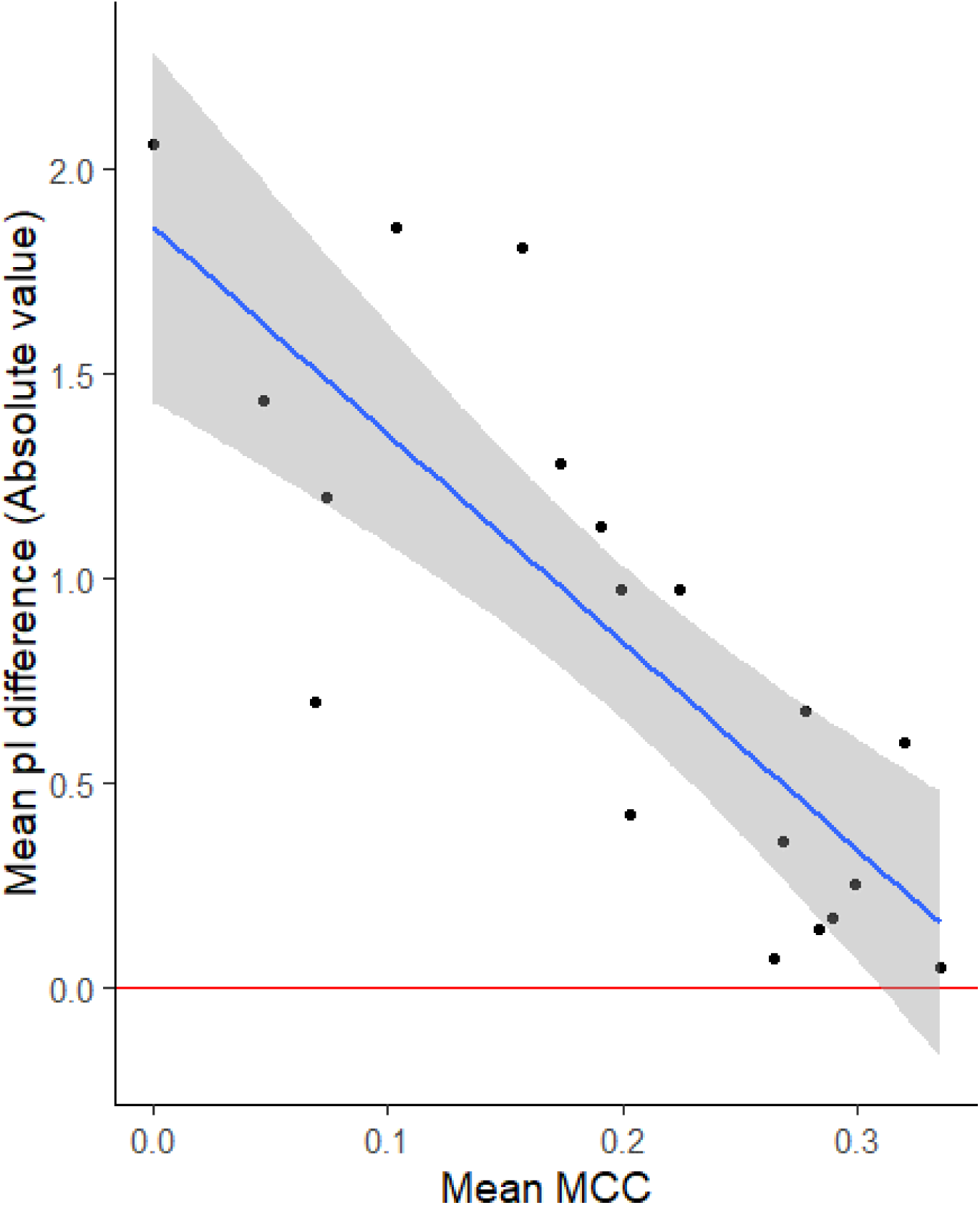
Correlation between polynucleotide-binding region prediction (represented as mean Matthews Correlation Coefficient, “Mean MCC” on the x axis) and pI prediction (“Mean pI difference” on the y axis). Mean MCCs were calculated for all polynucleotide-binding prediction methods evaluated for this study. The blue line indicates the least-squares linear regression, with a shaded 95% confidence interval.

The impact of other conserved structure predictions on pI largely aligned with known PBRs, as shown in Figure 7. This confirmation of structure-based pI prediction among closely-related viruses further supports the case for excluding PBRs from pI calculation. *Allolevivirus* phages Qβ (EBQB) and SP (EBSP), as well as *Tymovirus* turnip yellow mosaic virus (TYMV), all had prominent known PBRs located on beta turns. Accordingly, the beta turn prediction showed some of the greatest improvements in pI prediction for EBQB, EBSP, and TYMV. Predictions for other Tymoviruses improved as well, including belladonna mottle virus (BDMV), Erysimum latent virus (ELV), and Scrophularia mottle virus (ScrMV). Excluding beta sheets (independent of beta turns) also improved pI prediction for *Leviviridae* (EBFR, EBGA, EBMS2, EBQB, and EBSP) and Tymoviruses (ELV and ScrMV), as expected from known PBRs. However, neither beta sheet nor beta turn prediction performed as well overall as arginine-rich region prediction, even for *Leviviridae*.

**Figure 7:**
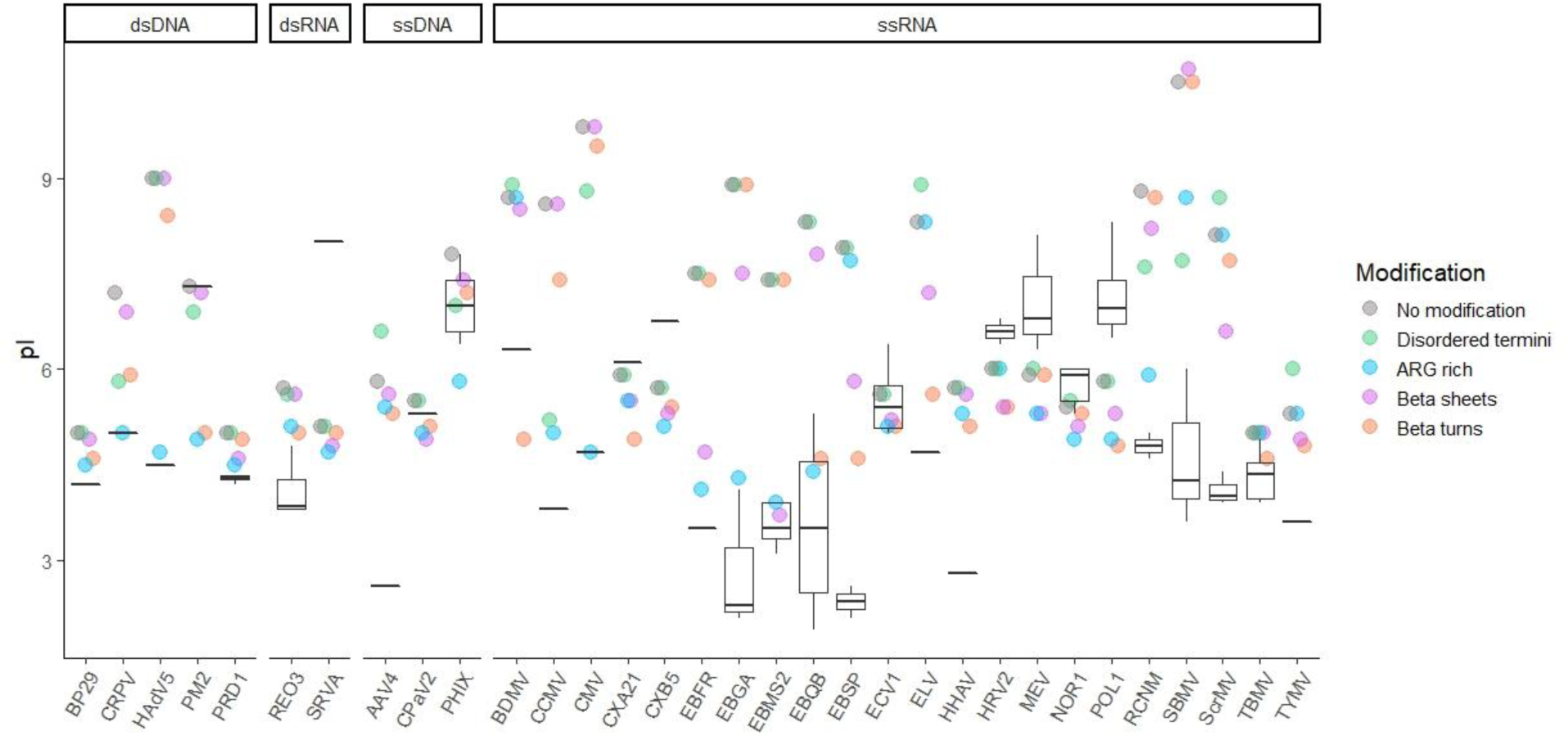
Effect of excluding particular types of predicted polynucleotide-binding regions (PBRs) on theoretical capsid pI. Box and whisker plots represent the range of empirical pIs found in the literature, while circles represent predicted pIs calculated without modification, as well as after excluding predicted PBRs. Several structures were used to predict PBRs: disordered N- and C-termini (“Disordered termini”), arginine-rich regions (“ARG rich”), beta sheets (“Beta sheets”), and beta turns (*i.e*., turns between adjacent beta sheets, “Beta turns”). A key to the 32 virus abbreviations (x axis) is provided in Table 2.

Several viruses had known PBRs on disordered N- or C-termini: cowpea chlorotic mosaic virus (CCMV), cucumber mosaic virus (CMV), cottontail rabbit polyomavirus (CRPV), red clover necrotic mosaic virus (RCNM), and southern bean mosaic virus (SBMV). These viruses were among the very few to show improved pI after removal of disordered termini, as shown in Figure 7. The specificity of pI improvement on removal of disordered termini both verifies the connection between PBRs and pI, and also indicates that the occurrence of disordered termini is not particular to PBRs and therefore a poor predictor.

## 3. Discussion

This study proposed a heuristic approach to pI prediction, in which only capsid residues bound to the viral polynucleotide were selectively excluded from the capsid charge calculation. This approach was based on the hypothesis that the charges of ionizable amino acids in these regions are neutralized by coulombic interactions with the virus polynucleotide. The PBR exclusion approach proposes a conceptual and conditional model for virion charge distribution, rather than a simplistic universal model. Furthermore, this approach excludes capsid regions only on the basis of polynucleotide binding, rather than for a desired impact on pI, and therefore avoids arbitrary removal of capsid regions to fit empirical pI values. A nonarbitrary approach is especially important given the relatively few viruses for which empirical pIs and capsid structures are known. With such a small data set, arbitrary variables are likely to overfit the model to the training set. In addition, the variety of capsid structures even among nonenveloped, icosahedral viruses may defy a universal model.

For example, bacteriophage MS2 has often been the exemplar of the effect of the viral genome on pI. The predicted pI of MS2 based on charged capsid moieties has been estimated between pH 7 (8) and 9 (51) (in this study, 7.4), while the estimated pI of MS2’s single-stranded RNA genome is approximately 3 (51). By contrast, the measured pI of MS2 is approximately 3.5 (7), closer to the estimated RNA pI than the estimated capsid pI. Structural models both including and excluding inner core charges have been proposed to explain this discrepancy. In MS2, the outermost shell of ssRNA lies directly beneath the capsid, and much of the capsid interior is devoted to binding the ssRNA genome (approximately 57% of the MS2 capsid protein) (32, 52). This highly basic region can be negated, thus decreasing the calculated pI to near the empirical value, by supposing either a strong negative influence from the core or that only exterior residues are relevant. In this way, bacteriophage MS2 is also an exemplar of the danger of overfitting a model.

### 3.1. Modifications based on known capsid structure

The first evidence in support of the PBR exclusion hypothesis came from 3D capsid structures. While exterior surface residues were not correlated with overall virion charge, removing interior capsid surfaces improved overall pI prediction for a diverse set of viruses (Figure 2). The exclusion of the interior capsid residues was most beneficial for viruses with interior PBRs. These viruses (*Leviviridae* phages and viruses with disordered termini) also show some of the greatest discrepancies between empirical and predicted pIs (Figure 3). Because these typically basic, arginine-rich PBRs stabilize the negatively-charged polynucleotide via coulombic forces (35, 36, 39, 41, 43), PBRs should therefore contribute no net charge to the virion. The degree to which major capsid proteins are involved in polynucleotide binding differs greatly between viruses, and is generally greatest for viruses with single-stranded genomes (32, 33, 36, 52). This tendency is corroborated by the greater effect of both PBR exclusion and interior capsid surface exclusion on pI. As shown in Figures 3 and 4, the one double-stranded virus showing a major change (>2 pH units) in predicted pI after PBR exclusion is human adenovirus 5 (HAdV5). The PBRs on HAdV5 are located on core proteins, however, rather than on the capsid shell. This variable response based on genome also indicates that a universal approach of removing the capsid interior may not yield the best overall fit to empirical pI values. Nonetheless, the results in Figure 3 demonstrate that removing the capsid interior may have negligible effect on viruses with evenly-distributed ionizable amino acids.

Following these insights from 3D capsid structures, known PBRs were identified for exclusion from the capsid charge calculation, based on reports from the literature or conserved PBR structures. Known PBRs were found for 15 viruses. Predicted pI values were calculated for both the unmodified proteomes of these viruses, as well as the proteomes after PBR exclusion. Exclusion of known PBRs yielded additional improvements in predicted pIs, especially for viruses with unavailable 3D structures (Figure 3). PBRs with the greatest impact on capsid pI fell into three broad structural categories: interior beta sheets, turns between beta sheets, and disordered termini. In addition, PBRs tended to have a high fraction of arginine and lysine (0.42 ± 0.29 arginine and/or lysine), simply termed “arginine-rich” here and elsewhere in the literature. These similarities indicated that PBRs could potentially be predicted via conserved primary and/or secondary structures.

### 3.2. Selective exclusion of predicted polynucleotide-binding regions

Based on the improvement in predicted pIs after excluding known PBRs, we attempted to predict PBRs based on these conserved structures. These predicted PBRs would then be excluded from predicted pI calculations. The structure-based PBR prediction method performed better than existing RNA- and DNA-binding prediction tools. Selection of arginine-rich regions was the most comprehensive predictor (MCC = 0.32 ± 0.29), although this prediction also improved by selecting basic beta turns (MCC = 0.34 ± 0.28). (The MCC values for all predictions are provided in Table S1.) Unlike disordered termini and many basic beta sheets, beta turns may be adjacent in tertiary structure and constitute a region of basic charges, while still being distant in the protein sequence. Therefore, many beta turn PBRs did not satisfy the conditions of arginine-rich prediction.

Two of the most conclusive findings from the predictive models were that A) pI prediction improved most for groups of viruses known to have certain conserved PBR structures, and B) better PBR prediction led to better pI prediction. When comparing pI predictions based on individual PBR structure predictions (Figure 7), pIs improved both for viruses known to have those PBRs, as well as closely related viruses. For example, Leviviruses and Tymoviruses, which feature PBRs along beta sheets and beta turns (53–55), showed the greatest pI improvements from beta sheet and turn predictions. Viruses with PBRs along disordered termini also showed the greatest pI improvement after exclusion of disordered termini. The strong correlation between PBR and pI prediction was true across the variety of prediction methods evaluated in this study, even though PBR prediction was conducted independently of the eventual impact on pI. Therefore, these PBR predictions further validate the results from known PBRs (Figure 3) to support the hypothesis that PBRs do not contribute to overall virion charge. Thus, unlike previous models for pI prediction (9, 11, 12, 17), the PBR exclusion method is a nonarbitrary method of predicting capsid pI, in that no part of the model was adjusted for the effect on pI.

While the PBR exclusion method explained many of the biggest discrepancies between empirical and predicted pIs, several exceptions indicate the need for further research and refinement. As shown in Figure 1, different strains of the same virus may deviate in pI by as much as 2 pH units. Of the 32 viruses considered here, seven predicted pI values remained more than 2 pH units from their empirical pIs after modification (Figure 5): adeno-associated virus 4 (AAV4), bacteriophage PM2 (PM2), simian rotavirus A (SRVA), bacteriophage SP (EBSP), poliovirus 1 (POL1), southern bean mosaic virus (SBMV), and Scrophularia mosaic virus (ScrMV). The pI prediction of EBSP, SBMV, and ScrMV all improved with modification, though further refinements may be needed to bring predictions closer to empirical pI values. Unfortunately, three of these viruses (AAV4, PM2, SRVA) are each represented by a single empirical pI reference, so the expected range of pIs for different strains and different experimental methods is unknown. Of these, both SRVA and PM2 are large viruses (Ø >75 nm) with multi-layered capsid structures, including an internal phospholipid bilayer in PM2. SRVA capsid proteins VP6 and VP7 also feature several calcium-binding regions (38). Polyvalent cations can be integral to the structure of some capsids, and may influence virion charge (56). However, whether integral cations contribute to overall charge differently than non-integral counterions in solution has yet to be determined.

Besides POL1, other members of the genus *Enterovirus* also had somewhat poorer pI prediction after removing predicted PBRs (Figure 4): coxsackievirus A21 (CXA21), coxsackievirus B5 (CXB5), echovirus 1 (ECV1), human rhinovirus 2 (HRV2), mengo encephalomyocarditis (MEV), and norovirus 1 (NOR1). The PBR prediction is likely to generate some false positives as well as false negatives. In a review of the literature, PBRs were only found for POL1, which has three interior, XNA-binding arginines (57). After removing these residues, the predicted pI of POL1 was slightly less accurate (Figure 3). By contrast, excluding the capsid interior slightly decreased the divergence between theoretical and empirical pIs for this group (Figure 3). Thus, *Enterovirus* may also share a similar capsid structure that was not explained by this attempt at PBR prediction. Hepatitis A virus (HHAV) is in the same family as Enteroviruses (*Picornaviridae*); yet, pI prediction for HHAV improved after excluding PBRs (Figure 4). However, unlike other Picornaviruses, HHAV occurs in cell culture and infected tissues in both enveloped and non-enveloped forms (26), and the sole source for HHAV pI is a brief with minimal information on methods (58). Therefore, further confirmation of empirical pIs for enveloped and non-enveloped HHAV is a research priority.

Picornaviruses (including Enteroviruses) have covalently-bound genomes (26), and may rely less on electrostatic binding than other ssRNA viruses. During assembly, Picornaviruses form procapsids. Though there is disagreement over the precise mechanism of encapsidation, the procapsid contains the ssRNA polynucleotide, and the mature virion is condensed around the core via cleavage and restructuring of capsid proteins (26). However, the interaction that stabilizes the virion is likely between capsid proteins and proteins involved in genome replication, *i.e*., a protein-protein interaction rather than a protein-RNA interaction (59). This alternate mechanism of stabilizing the ssRNA capsid may explain the poor performance of the PBR exclusion model for Enteroviruses in particular. However, this exception supports the fundamental hypothesis that polynucleotide-binding is responsible for the greatest pI discrepancies; for viruses known to lack PBRs, the PBR exclusion method is neither necessary nor appropriate. The more that is known about the diversity of virion morphogenesis, the more detailed our model of virion charge structure can become.

Michen and Graule (7) noted that different methods of measuring pI may be responsible for some of the variation in empirical pIs reported in the literature. The available empirical pIs for these enteroviruses were all determined via isoelectric focusing, However, extrapolation of pI from electrophoretic mobility is generally considered a more accurate method for determining pI of monodispersed viral particles (56). Virus aggregation in isoelectric focusing may also lead to inaccurate estimation of pI, since aggregates can be subject to gravitational/buoyant forces in addition to electrophoresis (60). However, observation of electrophoretic mobility via dynamic light scattering requires high titers (> 10^9^ PFU/mL) not reasonable for many viruses. For the viruses referenced in this study, empirical pIs determined via electrophoretic mobility and isoelectric focusing methods tended to agree for the few viruses tested using both methods, as shown in Figure S2. Isoelectric focusing was used for all of the high pI values (above pH 7) included in this study, while electrophoretic mobility measurements skew toward lower pIs. Since many of the high virus pIs have not been confirmed via multiple methods (including those of *Enterovirus*, as well as AAV4, PM2 and SRVA), further validation of empirical pIs is needed.

### 3.3. Future research needs

The PBR exclusion approach outlined in this study demonstrated strong potential to predict virus pI based on structural features. The insight of PBR exclusion provides researchers three avenues for predicting the pI of a virus, in order of preference: A) if the virus has a well-defined capsid with known PBRs, the known PBRs may be excluded; B) PBRs may be predicted via the definitions of conserved structures (arginine-rich regions and beta turns) provided in this study, or C) if the virus has a well-defined 3D structure, excluding the interior residues may approximate PBR exclusion. Where information is limited, researchers should cross-confirm using multiple methods.

Using the PBR exclusion approach with either known or predicted PBRs successfully explained many of the largest discrepancies between theoretical and empirical pIs. Given the variation in empirical pIs reported for different virus strains (Figure 1), accounting for such major discrepancies may be the limit of a predictive model at this time. Though the structure-based prediction used in this study outperformed existing XNA-binding prediction tools, the main hurdle to a predictive pI model is that the available empirical pI and PBR data are poorly corroborated and over-represent certain virus groups (*e.g*., *Leviviridae)*. Further research is needed to identify the full extent of PBRs in virus capsid proteins, as well as to determine or corroborate empirical pIs for a broader range of viruses. In addition, the identification of beta turns used in this method relied on NetSurfP 2.0 to predict secondary structure (61). However, the conservation of structure in viral PBRs (*e.g*., interior beta sheets, disordered termini, and associated alpha helices) suggests that a sequence-based prediction tool specifically trained to viral capsid PBR motifs could be successful in identifying PBRs directly from proteome sequences.

## 4. Methods

### 4.1. Prediction of isoelectric point

The sum of ionizable amino acid charges (Q) for a given capsid protein, *i*, was calculated using the Henderson-Hasselbalch formula:

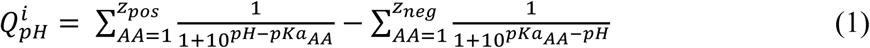

where *pKa*_*AA*_ is the pKa value for a given amino acid, *z*_*pos*_ is the number of positively charged amino acids in the capsid, and *z*_*neg*_ is the number of negatively charged amino acids in the capsid. Values for amino acid pKas are given in Table 1. The sum of charges for the entire capsid was considered by multiplying the sum of charges (including C- and N-termini) for each of *m* capsid proteins by the number of copies, *n*, of that protein within the capsid:

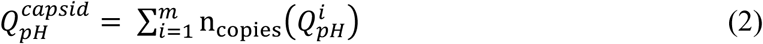

A summary of capsid proteins and copy numbers is presented in the Supporting Information (Table S3). Amino acids identified for exclusion (whether based on 3D capsid structure or PBRs) were not included in the sum of charges, including terminal amine and carboxyl charges where appropriate. Values of the predicted pI were calculated using the sum of charges for each capsid from pH 0 to 14 in order to find the pH value at which the charge approximated 0.

Empirical values for virus pIs were gathered from the literature, including Michen and Graule’s previous review (7), as well as all primary articles for additional information on experimental methods and conditions. A summary of empirical pIs is provided in the Supporting Information (Table S2). Multiple pIs have been reported by several sources using isoelectric focusing for mengovirus, human coxsackievirus A and B, human echovirus 1, and poliovirus 1 and 2 (7). While initially researchers suggested the different pIs represented different viral forms (62), Vrijsen et al. (60) demonstrated that the fraction of poliovirus 1 appearing near pH 4-5 in the pH gradient was separable by low centrifugation and did not appear when the virus sample was added near the primary pI (∼ pH 7) in an established pH gradient. Therefore, the secondary pI for poliovirus 1 was likely due to aggregated virus, while the primary pI represented monodispersed virus (60). Since the secondary band did not form when polioviruses were added at the pI (where some virus aggregation would also likely occur), the ampholytes used in isoelectric focusing as charge carriers may have destabilized viruses by charge neutralization and/or hydrophobic interactions. Ampholines used for isoelectric focusing have been shown to aggregate with acidic polysaccharides around pH 5 (63). Human enteroviruses B and C and mengovirus have also shown two distinct bands in isoelectric focusing with a secondary pIs at pH 4.4 – 4.8 (7). However, Chlumecka et al. (64) later found that the lower pI of mengovirus could be eliminated by adding ethylene glycol to promote dispersion. Here, all secondary pIs near pH 4-5 are likewise assumed to be an artifact of isoelectric focusing due to complexation with the ampholine buffer.

Our method of excluding polynucleotide-binding amino acids from the predicted pI calculation was evaluated by comparing the difference between predicted pIs and mean empirical values for each virus. The mean and standard deviation of these differences was compared before and after modifying the protein sequences by removing PBRs. The R stats package was used to perform a paired, two-tailed t-test between the modified and unmodified samples.

### 4.2. Sources of capsid structures

Amino acid sequences for all proteins composing the virus capsids were accessed via the UniProt database, as well as information about protein copy number, location within the capsid, and PBRs within capsid proteins, except as noted in the Supporting Information (Table S3) (38). Sequences were analyzed in FASTA format using scripts written in-house using the R language (65). Only regions known to bind to the viral genome were considered, not regions binding host ATP/nucleotides. For viruses in the family *Leviviridae*, the entire interior-facing beta sheet was considered polynucleotide-binding, as this region and RNA-associated assembly is highly conserved (31, 34). The beta sheet region was identified via literature values for bacteriophages MS2 (EBMS2) and Qβ (EBQB) (31, 52). The beta sheets for bacteriophages fr and GA were identified by cross-confirming results of visualization via PyMol (66), secondary structure prediction via NetSurfP 2.0 (61, 67), and sequence similarity via the UniProt database. For human adenovirus C5 (HAdV5), two of the proteins in the UniProt database were entirely located within the virion core and closely associated with the host genome: histone-like nucleoprotein (P68951) and core protein X (Q2KS10). The entire sequence of these two short proteins (173 and 19 amino acids, respectively) was considered polynucleotide-binding.

Viruses with available 3D capsid structures were identified via the VIPERdb icosahedral virus capsid database (68). The impact of 3D structure on capsid pI was evaluated by defining exterior residues as: A) including only the exterior capsid surface, or B) excluding only the interior capsid surface. Residues on the interior and exterior capsid surfaces were identified using the CapsidMaps tool via the VIPERdb website (30). In this study, 27 viruses were initially identified for having available 3D structures as well as empirical isoelectric point values in the literature. Of these 27 viruses, the surface residues of the 7 largest viruses (BP29, EBT4, HAdV5, PM2, PRD1, REO3, and SRVA) could not be accessed via the CapsidMaps tool due to large size and/or insufficient detail. A summary of viruses evaluated in this study is provided in Table 2. 3D structures for closely related viruses were used when complete structures for strains used in empirical pI measurements were not available: feline parvovirus (PDBID #1c8g) was used for canine parvovirus (CPAV2, 98.7% capsid protein sequence identity), human coxsackievirus B3 (PDBID #1cov) was used for human coxsackievirus B5 (CXB5, 90.4% genome polyprotein sequence identity), and human papillomavirus (PDBID #5keq) was used for cottontail papillomavirus (CRPV, 43.4% capsid protein sequence identity). Despite the relatively poor sequence similarity between the human and cottontail papillomavirus, both had similar pIs for both unmodified (7.25 ± 0.05) and (known) PBR-excluded (6.15 ± 0.15) proteome sequences. Papillomaviruses share a DNA-binding C-terminus on the L1 major capsid protein, and may also have a less essential PBR on the N-terminus of the L2 minor capsid protein (69, 70).

### 4.3. Predictive methods

Potential PBRs were identified based on proteome sequences alone in an attempt to predict virion pI by the PBR exclusion method for a group of 32 viruses. PBRs were first predicted by identifying conserved PBR structures, including: predominately basic beta sheets and associated beta turns, disordered C- and N-termini, and arginine- and lysine-rich regions. Prediction of secondary structures (beta sheets, beta turns, and disordered termini) from proteome sequences was performed using a deep-learning protein structure prediction tool, NetSurfP 2.0 (61, 67). In addition to identifying PBRs via conserved structures, two web-based tools for position-specific prediction of XNA-binding by residues were evaluated: Pprint for RNA-binding (45, 46, 50), and DRNApred (47, 48) for DNA- and RNA-binding regions (48). However, neither tool was developed for viral polynucleotide binding prediction specifically.

All PBR predictions were evaluated based on the mean Matthews Correlation Coefficient (MCC) for all proteins in the training or validation set (49):

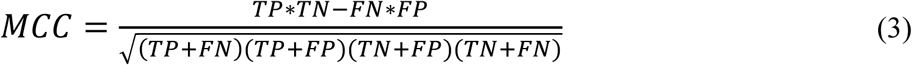

where TP (true positives) are correctly predicted residues within PBRs, FP (false positives) are incorrectly predicted residues outside of known PBRs, and FN (false negatives) are residues within known PBRs not predicted by the function. The MCC returns a value between -1 and 1, and has the benefit of evaluating predictions of both positive and negative values. Capsid proteins containing known PBRs for the viruses used in this study were used as a training set for identifying PBRs. Predictions were then evaluated against a validation set of 40 different viral capsid proteins selected from the UniProt database which contained DNA- or RNA-binding regions (38). A summary of capsid proteins in the training and validation sets is presented in the Supporting Information (Table S4). For each prediction method, the mean MCC for all viruses was compared to the mean absolute difference between theoretical and empirical pI for all viruses. The correlation between MCC and absolute pI difference was calculated via Spearman’s rank correlation using the R stats package (65).

Methods for predicting arginine-rich regions, beta sheets and turns, disordered termini, and RNA-binding via Pprint were optimized based on known PBR sequences. Variables taking an integer value (*e.g*., amino acid counts or distances along a sequence) were optimized via brute force calculation, while non-integer variables were optimized using a 1-dimensional optimization function, optimize, in the R stats package (65). Via optimization, arginine-rich regions were defined as regions consisting of at least 24% arginine and/or lysine, with a minimum of 5 arginines/lysines and a maximum separation of 9 amino acids between consecutive arginines/lysines. Though arginine was the predominate amino acid in these regions (thus the name “arginine-rich”), lysine also has a strongly basic side-chain and was frequently present in the same regions.

Beta sheets were defined as regions of contiguous predicted beta sheet structure at least 12 amino acids in length and composed of net neutral or basic ionizable amino acids (*i.e*., at least as many basic ionizable amino acids [ARG, LYS, HIS] as strong acidic ionizable amino acids [ASP, GLU, CYS]). Prediction of beta turn PBRs was optimized by searching for contiguous regions including a predicted beta turn and containing at least 2 basic ionizable amino acids (ARG, LYS, HIS) separated by at most 3 amino acids. Disordered termini were defined via optimization as regions including the C- or N-terminus containing only residues with a disorder probability of 0.66 or greater, based on randomness predictions by NetSurfP 2.0 (61, 67).

The optimal RNA-binding likelihood for Pprint (46), based on the support vector machine (SVM) score (Pprint’s output), was determined to be 0.57. Unlike Pprint, DRNApred classified residues using default parameters, so no optimization was required (47). However, the DRNApred prediction was better when a positive hit for either DNA- or RNA-binding was considered positive (mean MCC: 0.17 ± 0.23), than when only hits matching the virus genome type were considered positive (mean MCC: 0.10 ± 0.24). The poor classification of viral capsid PBRs based on nucleic acid type may result from DRNApred not being intended for use specifically with viral genomes, which may be single- or double-stranded DNA or RNA.

### 4.4. Data availability

All 3D structures, proteome sequences, and empirical pI data used in this study are publicly available. National Center for Biotechnology Information (NCBI) and Protein Data Bank (PDB) identifiers and citations for the viruses referenced in this study are provided in Table 2. A detailed summary of all capsid protein sequences, as well as UniProt entries and citations, is provided in Table S3. Secondary structures for proteome sequences were predicted via the NetSurfP-2.0 webtool (61, 67). PBRs were predicted using both in-house code to identify conserved structures based on NetSurfP output, as well as freely available webtools: Pprint (46) and DRNApred (47). In-house R scripts used to identify conserved PBR structures are provided in the Supporting Information, Section S5. Empirical pI data and citations are provided in Table S2.

